# Structural basis of substrate recognition and translocation by human ABCD1

**DOI:** 10.1101/2021.09.24.461565

**Authors:** Zhi-Peng Chen, Da Xu, Liang Wang, Cong-Zhao Zhou, Wen-Tao Hou, Yuxing Chen

## Abstract

Human ATP-binding cassette (ABC) transporter ABCD1 transports CoA esters of saturated/monounsaturated very long chain fatty acid from cytosol to the peroxisome for β-oxidation. Dysfunction of human ABCD1 usually causes the severe progressive genetic disorder X-linked adrenoleukodystrophy, which eventually affects the adrenal glands and/or the central nervous system. Here, we report three cryo-EM structures of human ABCD1 in various states. The apo-form ABCD1 at 3.53 Å resolution adopts an inward-facing conformation, harboring a phosphatidyl ethanolamine (PE) molecule at each lateral entry of substrate cavity. In the substrate-bound ABCD1 structure at 3.59 Å resolution, two molecules of C22:0-CoA (one of the physiological substrates of ABCD1) is symmetrically bound to the transmembrane domains (TMDs). Each C22:0-CoA adopts an unpresented L-shape configuration: the CoA portion inserts into a polar pocket at the TMD at a pose parallel to the membrane plane, whereas the acyl chain portion perpendicular to membrane plane is embedded in a hydrophobic pocket at the opposite TMD. Upon binding the two C22:0-CoA molecules, which resemble a pair of hinges crossing the two TMDs, the two nucleotide-binding domains (NBDs) of ABCD1 approach towards each other. Addition ATP to the substrate-bound ABCD1 enabled us to reveal an ATP-bound structure at 2.79 Å, which shows an outward-facing conformation with the dimerized NBDs succeeding substrate release. These three structures combined with biochemical assays exhibit a snapshot of ABCD1-mediated substrate recognition, translocation and release. These findings provide the structural insights into the transport mechanism of ABC transporters that transport amphipathic molecules with long acyl chains.

## Introduction

ATP-binding cassette (ABC) transporters are ubiquitous transmembrane proteins found in all living organisms. They utilize the energy from ATP hydrolysis to drive the transport of diverse substrates across membranes^1,2^. The 48 members of human ABC transporter superfamily are grouped into seven distinct subfamilies (ABCA to ABCG). The ABCD subfamily consists of four members: ABCD1, also known as adrenoleukodystrophy protein (ALDP), ABCD2 (ALDP related protein, ALDRP), ABCD3, also known as 70-kDa peroxisomal membrane protein (PMP70), and ABCD4 (PMP70-related protein, P70R)^3–6^. As half transporters, ABCD transporters are expressed as single polypeptides containing one transmembrane domain (TMD) and one nucleotide-binding domain (NBD). They usually function as homodimers^7–9^, although heterodimers of ABCD1~3 have been observed in some studies^10,11^. The previously reported structure of ABCD4 revealed a distinct substrate and function from that of ABCD1~3^12^. ABCD1~3, which are localized in the peroxisomal membrane^6,13,14^, transport acyl-CoA esters into the peroxisome for β-oxidation^8,15–17^. In contrast, the putative cobalamin transporter ABCD4 is localized in the membrane of lysosome^18,19^.

Although ABCD1~3 all transport acyl-CoA esters, they differ from each other in substrate specificity. ABCD1 prefers to transport CoA esters of saturated and monounsaturated very long chain fatty acid (VLCFA-CoAs), such as C22:1-CoA, C22:0-CoA and C26:0-CoA^8,20^. ABCD2, which shares a sequence identity of 62% with ABCD1, is capable of transporting most substrates of ABCD1, in addition to CoA esters of polyunsaturated fatty acids (PUFA-CoAs) such as C22:6-CoA and C24:6-CoA^11,20,21^. However, ABCD3, which is 39% sequence-identical to ABCD1, specifically transports CoA esters of dicarboxylic acid, branched-chain fatty acid, and the bile acid intermediates di- and tri-hydroxycholestanoic acid (DHCA and THCA)^16,22^. These acyl-CoA esters transported to the peroxisome by ABCD1~3 are subject to the β-oxidation of fatty acids or biosynthesis of bile acids. Thus, dysfunctions of these transporters usually cause severe diseases^17^.

Defects of ABCD1 cause X-linked adrenoleukodystrophy (X-ALD) due to the failures of VLCFAs transportation into peroxisome^4,15^. X-ALD is a severe progressive genetic disorder affecting the adrenal glands, the spinal cord, and the white matter (myelin) of the nervous system, which occurs in 1/20,000 males. Among a wide range of phenotypes of X-ALD, childhood cerebral adrenoleukodystrophy (CCER) is the most common phenotype with onset of neurological symptoms in children of 3~10 years old, which will rapidly develop to central nervous system demyelination, and eventually causes totally disability or even death within a few years^23,24^. Despite extensive biochemical and functional investigations on ABCD1 have been reported^15,25–27^, the molecular mechanism on how ABCD1 mediates acyl-CoA transport remains obscure.

Here, we report the structures of human ABCD1 determined by single-particle cryo-electron microscopy (cryo-EM) in three distinct states: the apo form at 3.53 Å, the substrate- and ATP-bound structures at 3.59 and 2.79 Å, respectively. Combined with the biochemical assays, these structures revealed a snapshot of the transport cycle of ABCD1, and provided hints for the substrate recognition and transport mechanism of the acyl-CoA esters transporters.

## Results

### Biochemical characterization and structure determination of human ABCD1

We initially overexpressed the full-length human ABCD1 (hABCD1) in HEK293F cells, but it is not sufficient for further study due to the very low expression level. In contrast, the expression level of *Caenorhabditis elegans* PMP-4, a 52% sequence-identical homolog of hABCD1, is much higher. Sequence alignment (Extended Data Fig. 1a) indicated that hABCD1 and PMP-4 differ from each other mainly in the N-terminal segment, which may be involved in the subcellular location^28^. Therefore, we constructed a chimeric version of ABCD1 (termed chABCD1), with the N-terminal 65 residues of PMP-4 joined to the core domains of human ABCD1. As expected, the expression level of chABCD1 is ~15 folds to that of hABCD1. The purified chABCD1 was homogenous in detergent micelles (Extended Data Fig. 1b). Both hABCD1 and chABCD1 proteins were extracted with lauryl maltose neopentyl glycol (LMNG) plus cholesteryl hemisuccinate (CHS) for further biochemical characterizations.

The ATPase activity assays of hABCD1 or chABCD1 in the presence of various VLCFA-CoAs or acetyl-CoA showed that the ATPases activity of either hABCD1 or chABCD1 could be stimulated by CoA esters of VLCFAs (Fig. 1b and Extended Data Fig. 1c,1d). Remarkably, chABCD1 displayed much higher activities than hABCD1 in the presence of VLCFAs-CoAs including C22:0-CoA, C24:0-CoA or C26:0-CoA. In contrast, addition of acety-CoA, or C26:0 and CoA separately (C26:0/CoA) displayed no change of activity. These results suggested that both hABCD1 and chABCD1 were at a physiologically relevant state, and VLCFAs-CoAs are their substrate, in agreement with previous reports^8,26,29^. We further measured substrate-stimulated ATPase kinetics parameters of chABCD1 (Fig. 1c). The results showed that C22:0-CoA, C24:0-CoA and C26:0-CoA displayed similar *K*_*m*_ values at ~2 μM and *V*_*max*_ of ~200 mol Pi/min/mol protein, suggesting similar specificities of chABCD1 toward these substrates.

**Fig. 1.**
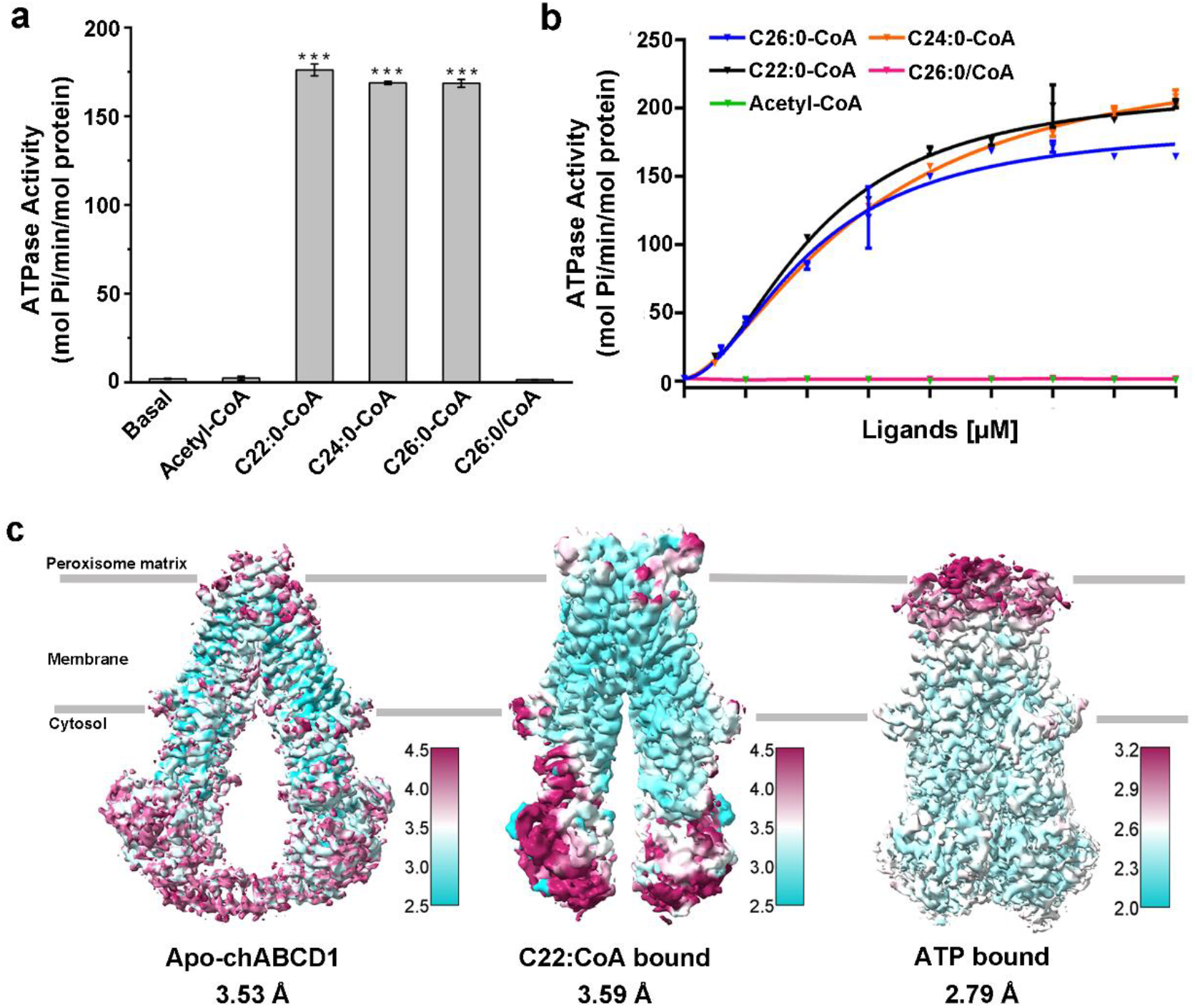
Function Characterization and Structure Determination of ABCD1. **a**, Stimulated ATPase activity of chimeric ABCD1 (chABCD1) over various CoA esters of very long chain fatty acid (VLCFA-CoAs) or acetyl-CoA. All data points analyzed above represent means of three independent measurements. Error bars indicate standard deviation. Unpaired two-sided t-test is used for the comparison of statistical significance. The P values of <0.05, 0.01, and 0.001 are indicated with *, ** and ***. **b**, Substrate-stimulated ATPase kinetics of chABCD1 over concentrations of VLCFA-CoAs or acetyl-CoA. **c**, Side views of 3D reconstruction of three ABCD1 structures. Density map colored by local resolution estimation in relion 3.1 or cryoSPARC 3.2.

To gain molecular insights into ABCD1, we used single particle cryo-EM to determine the structures chABCD1 in the detergent digitonin. In total, we solved three structures of chABCD1 (Fig. 1c): an apo-form chABCD1 at the resolution of 3.53 Å (Extended Data Fig. 2), a C22:0-CoA bound chABCD1 at 3.59 Å (Extended Data Fig. 3) and an ATP-bound chABCD1 at 2.79 Å (Extended Data Fig. 4). Notably, the N-terminal segment of PMP-4 is missing in all three structures, possibly due to its flexibility; the core domains of human ABCD1 could be clearly traced (termed ABCD1 hereafter).

### The apo-form structure of ABCD1

The overall structure of ABCD1 exhibits a two-fold symmetric homodimer consisting of two subunits, each of which contains a TMD and an NBD (Fig. 2a). The apo-form ABCD1 adopts an inward-facing conformation. Each TMD of ABCD1 consists of six transmembrane helices (TMs), which are tightly packed against each other at the peroxisome matrix side, but extend to the cytosol to form two diverged “wings”. The helices TM4 and TM5 from one TMD are swapped to the opposite TMD, which is a typical feature of type-IV ABC transporters^2^. Two pairs of coupling helices from the TMDs are embedded in the grooves on the NBDs, coupling the conformational changes between TMDs and NBDs. We found an extra density between TM5 and TM6 of each subunit, the shape of which is reminiscent of a lipid molecule (Fig. 2a,2b and Extended Data Fig. 2e). Accordingly, we fitted this density with a molecule of phosphatidyl ethanolamine (PE), which was probably introduced during protein purification. The PE molecule is located at the lateral entrance of the V-shape cavity of the TMD, and is accommodated by hydrophobic residues from TM5 and TM6 (Extended Data Fig. 5a).

**Fig. 2.**
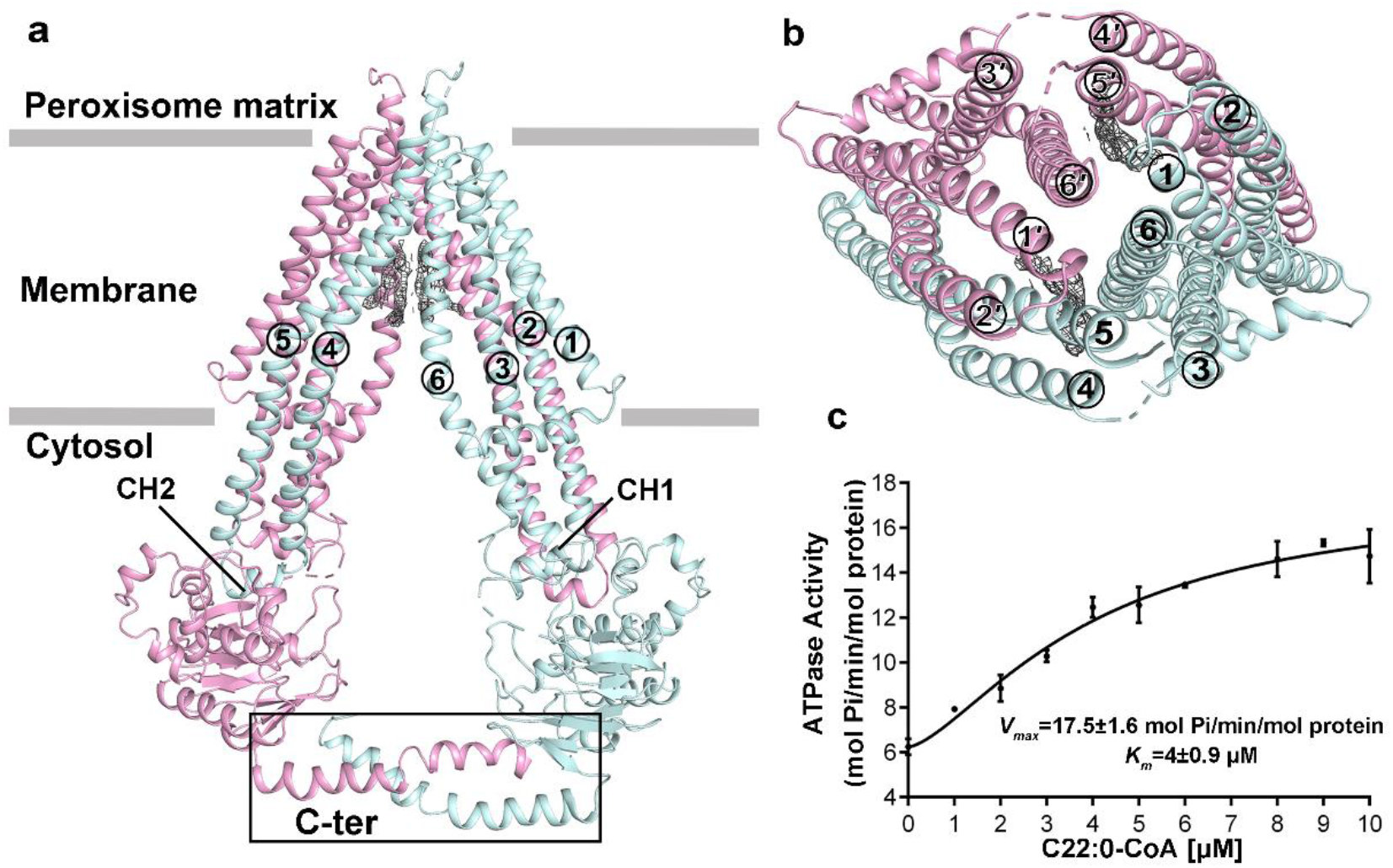
Overall structure of apo-form ABCD1. **a**, The structure of apo-form ABCD1 is shown in cartoon representation. The two monomers are colored in cyan and pink, respectively. Transmembrane helices (TMs), C-terminal coiled-coils and coupling helices (CH) are labeled. Two extra densities are shown in gray mesh. The peroxisome membrane plane is indicated as the gray rectangle. **b**, Top view of ABCD1 from the peroxisome matrix side. The six TMs of each monomer are numbered. **c**, The C22:0-CoA stimulated ATPase activity of the C-terminal coiled coils truncation variant of ABCD1.

The NBDs possess a classic NBD fold of ABC transporter in addition to a couple of helices that form a coiled coil at the C-terminus of the NBD (Fig. 2a). Sequence alignment revealed that this C-terminal coiled-coil helix is conserved in hABCD1 orthologs (Extended Data Fig. 5b). Truncation of this C-terminal helix led to a *K*_*m*_ value (4±0.9 μM) similar to the wild type, but a sharply decreased *V*_*max*_ (17.5±1.6 mol Pi/min/mg protein) (Fig. 2c). We thus speculated that this C-terminal coiled-coil may be important for stabilizing the two separated NBDs to maintain a conformation favored for ATP hydrolysis. In fact, a variant T693M on this C-terminal helix, which abolishes the function of ABCD1 without impacting the expression of ABCD1 in patients, is associated with X-ALD^30^.

### The substrate-bound structure revealed two portions of C22:0-CoA molecule respectively bind to the two TMDs of ABCD1

We obtained the complexed structure of substrate-bound ABCD1 by addition of 0.5 mM C22:0-CoA, which also adopts an inward-facing conformation, similar to the apo-form structure (Fig. 3a). However, the C-terminal coiled-coil helices in apo-form structure are missing upon substrate binding. Superposition of the C22:0-CoA-bound ABCD1 structure with the apo form yielded a root-mean-square-deviation (RMSD) of 9.20 Å over 1078 Cα atoms (Fig. 3b). Upon substrate binding, most TMs undergo a rigid body shift toward the 2-fold axis symmetry, and TM4 located in the transport cavity is split into two helices. These conformational changes of TMDs are transduced to NBDs, which approach to each other (Fig. 3c). Notably, a segment of residues 363-371, which is localized in the peroxisome matrix, folds into a short α-helix (the helix in the peroxisome, PH) in the C22:0-CoA-bound ABCD1 (Fig. 3a, 3b and Extended Data Fig. 3e). A similar helix was also found in a type-IV ABC transporter, *Campylobacter jejuni* lipid-linked oligosaccharide flippase PglK and its deletion abolished the substrate-stimulated ATPase activity^31^.

**Fig. 3.**
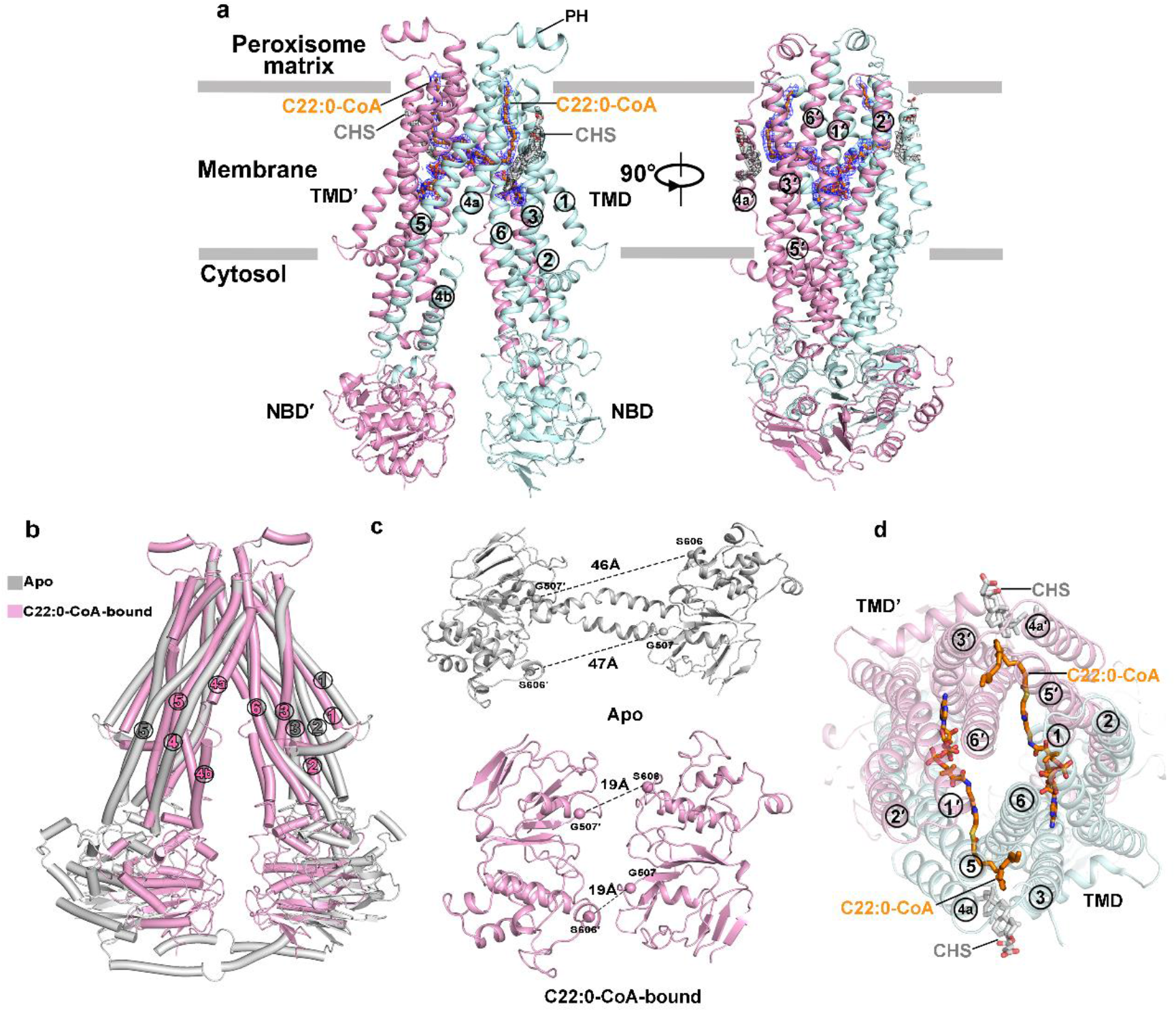
Overall structure of substrate-bound ABCD1. **a**, The structure of substrate-bound ABCD1 is shown in cartoon representation. The two monomers are colored in cyan and pink, respectively. Two pairs of extra densities are shown in gray mesh. The peroxisome membrane plane is indicated as the gray rectangle. The helices in the peroxisome matrix (PH) are labeled. **b**, Superposition of apo-form ABCD1 (grey) against C22:0-CoA-bound ABCD1 (pink). **c**, Distances between Walker A motif and signature motif of the two NBDs in the apo-form (grey) and C22:0-CoA-bound ABCD1(pink), respectively. **d**, Top view of C22:0-CoA-bound ABCD1 from the peroxisome matrix side. C22:0-CoA and cholesteryl hemisuccinate (CHS) molecules are fitted in the extra densities and shown as sticks.

Remarkably, a pair of L-shaped densities crossing the cavity of TMDs are symmetrically bound to ABCD1, each of which could be perfectly fitted with a molecule of C22:0-CoA (Fig. 3a, and Extended Data Fig. 3e). The acyl chain of C22:0-CoA inserts into the TMDs in parallel to surrounding TMs, whereas the portion of CoA lies in hydrophilic pocket in parallel to the membrane plane (Fig. 3d, 4a). The 3′-phospho-ADP moiety of the CoA is buried in a pocket formed mainly by a series of positively-charged residues (Fig. 4b). In detail, the adenine ring of the ADP is stabilized by Lys213 and Ser213 from TM3, and Arg401 from TM6 by hydrogen bonds, in addition to a π-cation interaction between Arg401 and the base. The 3′-phosphophate of ribose forms a salt bridge with Arg152 on TM2 and a hydrogen bond with Gln332′ from TM5′, whereas the diphosphate group interacts with Arg104 from TM1, Lys336′ and Tyr337′ from TM5′ by salt bridges and hydrogen bonds. The pantothenate and cysteamine moieties, which link the acyl chain and the 3′-phospho-ADP, extends from TMD to opposite TMD′ across the transport cavity (Fig. 3d, 4a), almost without direct interactions with TMDs except for the hydrophobic contact between Trp137 from TM2 with the pantothenate moiety (Fig. 4b). The 22-carbon acyl chain succeeding the pantothenate and cysteamine moieties is embedded in the TMD′. It takes a 90° turn at the sixth carbon and runs along the TM3′ and TM4′, up to the boundary of the peroxisome matrix. The acyl chain is lined in a hydrophobic cleft formed by a cluster of hydrophobic residues including: Val222′, Leu229′ and Ala233′ from TM3′, Pro243′, Val247′ and Val251′ from TM4′, Trp339′, Gly343′, Met346′, Val347′, Ile350′ and Ile351′ from TM5′, Ala384′, Ala388′, L392′, Ala395′ and Ala396′ from TM6′ (Fig. 4c).

**Fig. 4.**
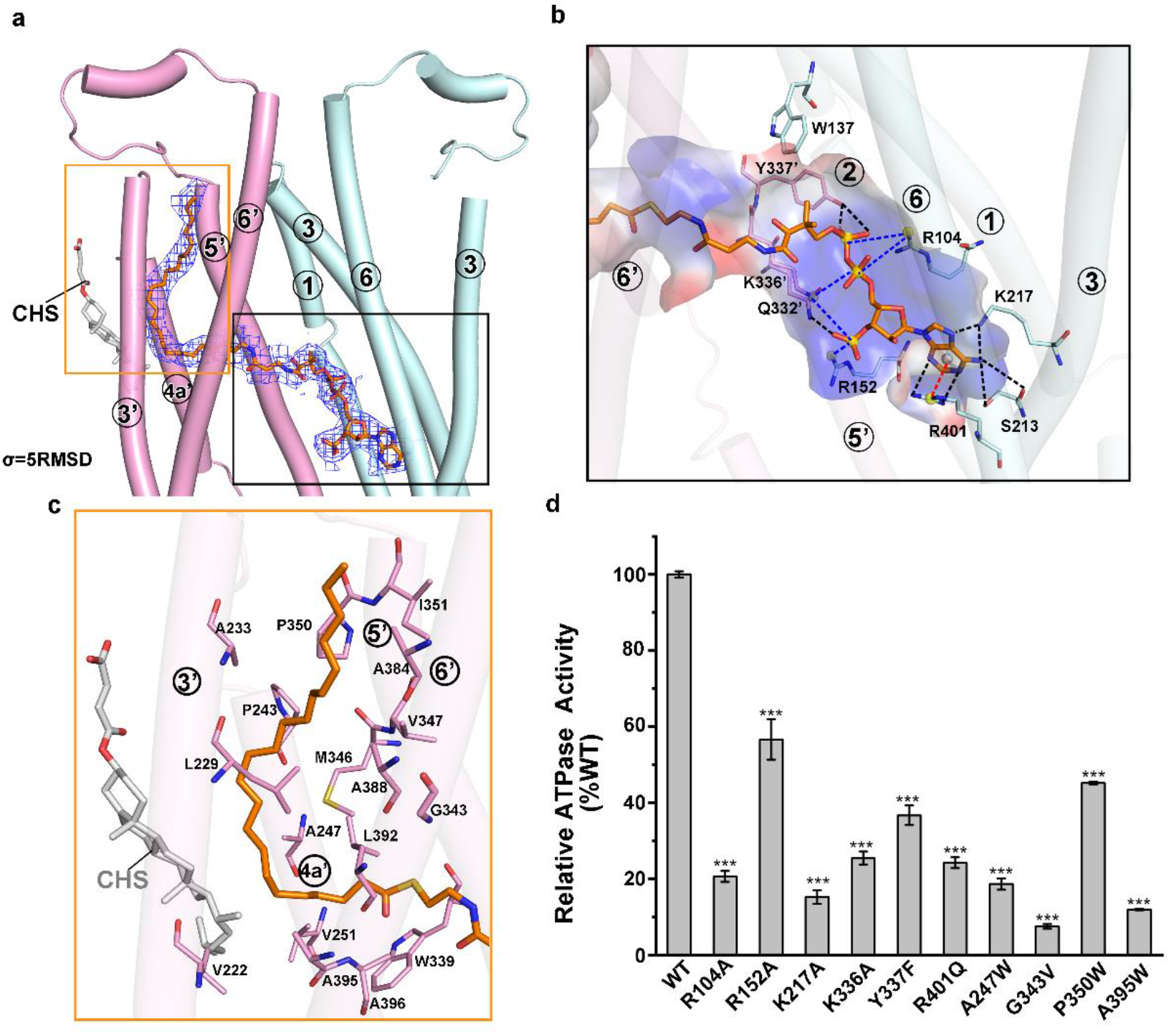
The C22:0-CoA binding sites of ABCD1. **a**, The two binding sites of the acyl chain and the CoA portion of one C22:0-CoA molecule. The two TMDs are colored in cyan and pink, respectively, and TMs are labeled for each TMD. **b**, The binding site of the CoA portion. Interacting residues are shown in sticks. Hydrogen bonds and salt bridges are indicated as the black dotted lines and blue dotted lines, respectively. One π-cation is indicated as the red dotted line. The electrostatic surface properties of the binding site are color-coded by electrostatic potential generated by PyMOL. **c**, The binding site of the acyl chain. Interacting residues are shown in sticks. **d**, C22:0-CoA-stimulated ATPase activity assays of mutants associated with substrate binding. At least three independent assays were performed for each assay. The means and standard deviations were calculated and the data are presented as means ± S.D. Error bars indicate standard deviation. Unpaired two-sided t-test is used for the comparison of statistical significance. The P values of <0.05, 0.01, and 0.001 are indicated with *, ** and **

At the peripheral of these two C22:0-CoA molecules, there are two extra densities that fit the detergents CHS introduced during protein purification (Fig .3a, 4c and Extended Data Fig. 3e). The CHS molecule, which inserts in the cleft between TM3 and TM4, provides an extra hydrophobic interface against the acyl chain (Fig. 4c). This CHS molecule might structurally mimic the cholesterol ester in membrane lipid.

We further generated variants with single mutations of substrate-binding residues in ABCD1 and performed the ATPase activity assays (Fig. 4d). The variants R104A, R252A, K217A, K336A, Y337F, and R401Q were designed to disrupt the polar interactions with CoA, and bulky sidechain substitutions A247W, G343V, P350W, A395W to the disrupt the hydrophobic interactions with the acyl chain displayed significant decrease of C22:0-CoA-stimulated ATPase activity compared to the wild type. Of note, R401Q and G343V were reported to be associated with X-ALD^30^. In sum, the extensive polar and hydrophobic interactions ensure a highly specific binding site toward the CoA and acyl chain portion, respectively. Sequence alignment revealed that these substrate-binding residues are highly conserved in ABCD1 homologs (Extended Data Fig. 6).

### The ATP-bound structure showed an outward-facing conformation of ABCD1

To investigate the mechanism of substrate release, we solved the ATP-bound structure of ABCD1 at a resolution of 2.79 Å (Extended Data Fig. 4) by addition ATP to the substrate-bound ABCD1. The E630Q mutant, which abolishes the ATP hydrolysis activity but maintains the ATP-binding capacity was introduced for ATP binding. Notably, two clear densities, which are sandwiched between the two NBDs, could be fitted with two ATP-Mg^2+^ molecules (Fig. 5a and Extended Data Fig. 4e). The structure showed excellent side-chain densities for most of the protein except that the helix PH and coiled-coil helices are missing. The ATP-bound structure adopts an outward-facing conformation (Fig. 5a), in which the two NBDs are dimerized and exhibit a typical “head-to-tail” configuration like all ABC transporters. The two ATP molecules bind between the Walker A motif of one NBD and the ABC signature motif of the opposite NBD. Despite 0.5 mM C22:0-CoA was added in prior to the addition of 20 mM ATP/Mg^2+^ during the cryo-EM sample preparation, the substrate C22:0-CoA is absent in the ATP-bound structure. Thus, the present outward-facing structure most likely represents a state succeeding substrate release upon ATP binding.

**Fig. 5.**
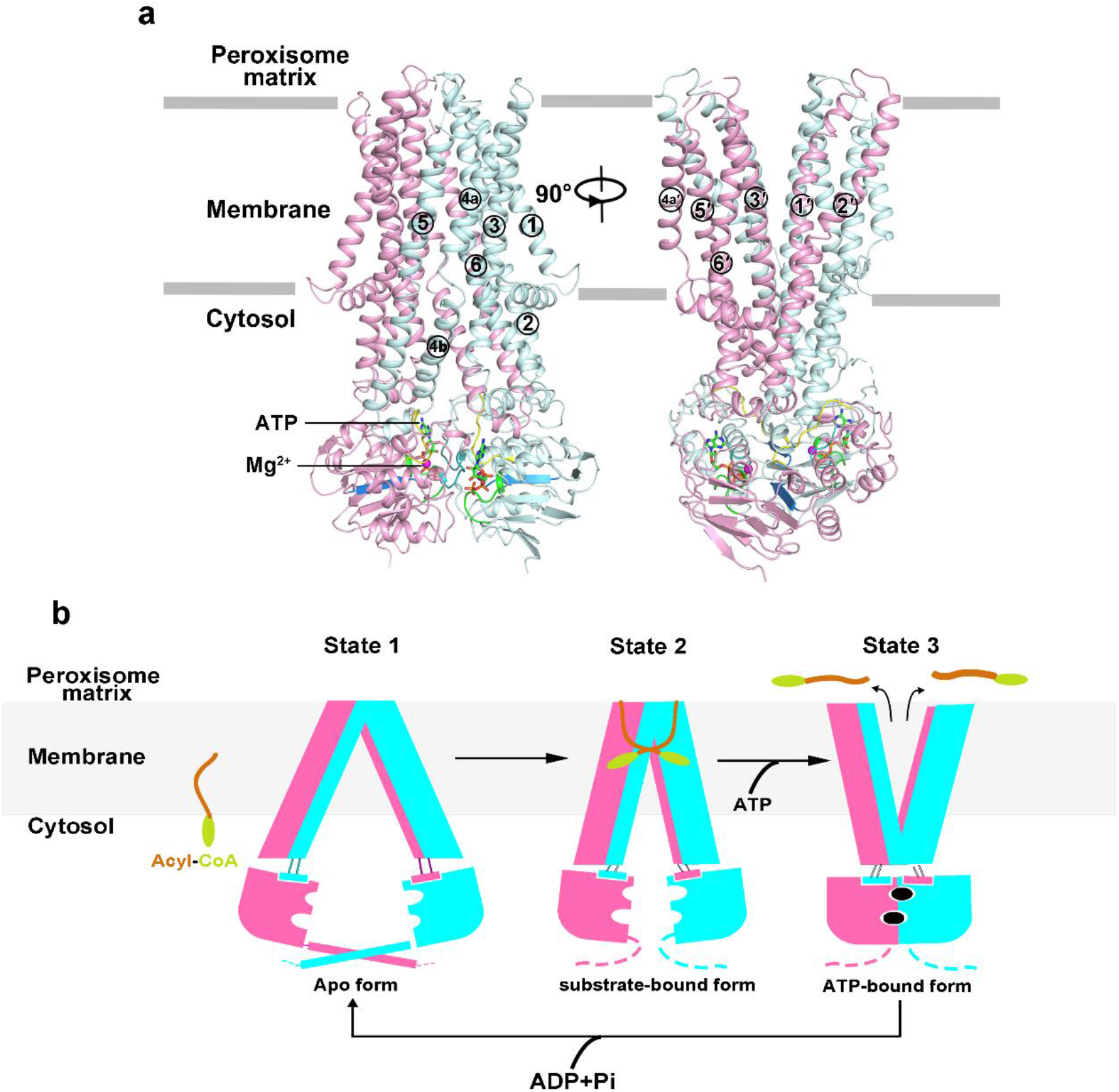
The ATP-bound ABCD1 structure and a proposed transport cycle of ABCD1 based on the three structures. **a**, The structure of ATP-bound ABCD1 is shown in cartoon representation. ATP/Mg^2+^ molecules are shown as sticks and balls. The Q-loop of NBD is colored in yellow, Walker A motif is colored in green, the signature motif is colored in teal, and Walker B motif is colored in marine. **b**, A model for ABCD1-mediated acyl-CoA translocation. The apo-form ABCD1 adopts an inward-facing conformation, which is further stabilized by the C-terminal coiled-coil helices (State 1). The substrate binding pulls two TMDs approach toward each other resulting in a narrower inward-facing transport cavity (State 2). Upon ATP binding, the dimerized NBDs make ABCD1 in an outward-facing conformation, ultimately facilitating the substrate release to the peroxisome matrix (State 3). Finally, the hydrolysis of ATP triggers a conformation change of ABCD1 to the rest state ready for another transport cycle.

## Discussion

The structure of C22:0-CoA-bound ABCD1 revealed a unique substrate binding pattern in which two C22:0-CoA is respectively bind to the two TMDs forming a L- shape conformation. For each C22:0-CoA, the CoA portion inserts into the hydrophilic transmembrane cavity in one TMD parallel with the membrane plane, while the acyl chain portion runs perpendicularly to the membrane plane, and extends to the opposite TMD, which provides a hydrophobic cleft for embedding the very long acyl chain. This binding pattern is distinct from other known-structures of ABC transporter that transports the amphipathic molecules with long acyl chains. For instance, the substrate-bound structures: lipopolysaccharide (LPS) flippase MsbA^32,33^, human phospholipid transporter ABCB4^34^ and phospholipid retrograde transport MlaFEDB^35^. All of these structures showed that the substrate is entirely located in the central cavity between the TMDs (Extended Data Fig. 7). Overall, our structure of C22:0-CoA-bound ABCD1 exhibit an unprecedented substrate-binding pattern for amphipathic substrates with long acyl chains.

The present structures of ABCD1 in the three different states combined with biochemical assays enabled us to propose a mechanism model of ABCD1-mediated VLCFA-CoA transport (Fig. 5b). In the apo-form state, ABCD1 adopts an inward-facing conformation, which is further stabilized by the C-terminal coiled-coil helices (State 1). The substrate binding pulls two TMDs approach toward each other resulting in a narrower inward-facing transport cavity (State 2). Upon ATP binding, the dimerized NBDs make ABCD1 in an outward-facing conformation, ultimately facilitating the substrate release to the peroxisome matrix (State 3). Finally, the hydrolysis of ATP triggers a conformation change of ABCD1 to the rest state ready for another transport cycle.

## Acknowledgments

We thank Dr. Yongxiang Gao at the Center for Integrative Imaging, Hefei National Laboratory for Physical Sciences at the Microscale, University of Science and Technology of China during cryo-EM image acquisition for ATP-bound ABCD1. We thank Xiaojun huang and Xujing Li for technical support on cryo-EM data collection for apo ABCD1 and C22:0-CoA-bound ABCD1 complex at the Center for Biological Imaging at the Institute of Biophysics (IBP), Chinese Academy of Sciences. This work was supported by the Ministry of Science and Technology of China (2020YFA0509302) and National Natural Science Foundation of China (32071206).

## Author Contributions

Y.C. and W.-T.H. conceived the project and planned the experiments. D.X., and Z.-P.C. expressed and purified human ABCD1. Z.-P.C. and L.W. performed cryo-EM data collection, structure determination and model refinement. Z.-P.C. performed functional assays. W.-T.H., Z.-P.C., Y.C. and C.-Z.Z. wrote the manuscript.

## Declaration of interests

The authors declare no competing interests.

## Data availability

The cryo-EM structures of apo-form ABCD1, C20:0-CoA-bound ABCD1, and ATP-bound ABCD1 have been deposited at PDB under the codes of xxxx, xxxx and xxxx, respectively. The cryo-EM density maps of three structures have been deposited at the Electron Microscopy Data Bank (EMD-xxxxx, EMD-xxxxx and EMD-xxxx, respectively).

## Methods

### Protein expression and purification

The codon-optimized full-length human *ABCD1* gene and *Caenorhabditis elegans* PMP-4 were synthesized by GENEWIZ Company, while the gene fragment encoding for chimeric protein was generated by overlap PCR during which a gene fragment encoding for residues 1-65 were amplified from *pmp4* and substituted the gene fragment encoding for residues 1-63 of human ABCD1. The wild type gene was cloned into a modified pCAG vector with an N-terminal FLAG tag (DYKDDDDK). Point mutations were introduced using a standard two-step PCR.

For protein expression, the HEK293F cells were cultured in SMM 293T-Ⅱ medium (Sino Biological Inc.) at 37°C, under 5% CO_2_ in a shaker. The cell transfection was performed when cell density reached ~2.5×10^6^ cells per mL. For 800 mL HEK 293F cell culture, ~2 mg plasmids premixed with 4 mg linear polyethylenimines (PEIs) (Polysciences, Inc) in 45 mL fresh medium for 15 min, then the mixture and another 150 mL fresh medium were added into the cell culture, followed by 30-min static incubation. The transfected cells were grown at 37°C for 12 hr, then 10 mM sodium butyrate (Aladdin) was added, and cultured at 30°C for additional 48 hr before being collected. After centrifugation at 4,000 g for 5 min, the cell pellets were resuspended in the lysis buffer containing 25 mM Tris-HCl pH 7.5, 150 mM NaCl, 20% glycerol (v/v), 1mM dithiothreitol (DTT) and 1×ethylene diamine tetraacetic acid (EDTA) free protease inhibitor cocktail (TargetMol). The suspension was frozen in liquid nitrogen and stored at −80°C for further use.

For cryo-EM protein sample preparation, membrane proteins were extracted from the cell with 1% (w/v) lauryl maltose neopentyl glycol (LMNG, Anatrace), 0.1% (w/v) cholesteryl hemisuccinate (CHS, Anatrace) and rotated gently at 4°C for 2 hr. After centrifugation at 45,000 rpm for 45 min (Beckman, Type 70 Ti), the supernatant was collected and applied to anti-FLAG M2 affinity gel (Sigma Aldrich) at 4°C for 1 hr. Then the resin was rinsed with wash buffer A containing 25 mM Tris-HCl pH 7.5, 150 mM NaCl, 10% glycerol (v/v),1 mM DTT, 0.06% digitonin (w/v). The protein was eluted with wash buffer B containing 25 mM Tris-HCl pH 7.5, 150 mM NaCl, 5% glycerol (v/v), 1mM DTT, 0.06% digitonin (w/v) supplemented with 200 μg/mL FLAG peptide. The protein eluent was concentrated by a 100-kDa cut-off Centricon (Millipore) and further purified by size-exclusion chromatography using a Superdex 200 Increase 10/300 (GE Healthcare) equilibrated with wash buffer C containing 25 mM Tris-HCl pH 7.5, 150 mM NaCl, 1mM DTT, 0.06% digitonin (w/v). Peak fractions were pooled and concentrated for biochemical studies or cryo-EM experiments.

The protein used for ATPase activity assay were also expressed and purified in the same way except that the detergent for wash buffer A, B, C were substituted by 0.005% LMNG and 0.0005% CHS.

All the mutated proteins using in this project were expressed and purified in the same way as the wild-type protein.

### ATPase activity assay

ATPase activities of wild-type human ABCD1, chimeric ABCD1 and its mutants were measured using the ATPase Colorimetric Assay Kit (Innova Biosciences) in 96-well plates at OD_630 nm_. All potential substrates including acetyl-CoA, behenoyl Coenzyme A (C22:0-CoA, ammonium salt), lignoceroyl Coenzyme A (C24:0-CoA, ammonium salt), hexacosanoyl Coenzyme A (C26:0-CoA, ammonium salt), Coenzyme A (CoA, trilithium salt) and Hexacosanoic acid (C26:0) were purchased from Sigma-Aldrich and solved in 5% (w/v) methyl-β-cyclodextrin (Sigma Aldrich).

To measure the ATPase activities of ABCD1 and chABCD1 upon different substrates, a final concentration of 0.03 μM protein was added to the reaction buffer containing 20 mM Tris-HCl, pH 7.5, 50 mM KCl, 1 mM DTT, 0.005% (w/v) LMNG/0.0005% (w/v) CHS, and 2 mM MgCl_2_ to 100 μL as one reaction sample. Then, potential substrates were diluted into different concentration and added into the reaction mixture. The mixture was incubated statically on the ice for 10 min and supplemented with 2 mM ATP before reactions were performed at 37°C for 20 min. then the amount of released Pi was quantitatively measured and statistical analysis was performed using Origin 2021b (Academic).

### Cryo-EM data collection

For the apo ABCD1 sample, two datasets with a total of 7495 micrograph stacks were automatically collected individually with SerialEM^36^ on a Titan Krios microscope at 300 kV equipped with a K3 Summit direct electron detector (Gatan) at a nominal magnification of 22,500× with defocus values from −2.0 to −1.5 μm. Each stack was exposed in super-resolution mode, resulting in 32 frames per stack, and the total dose for each stack was 60 e^−^/Å^2^. For the C22:0-CoA-bound ABCD1 samples, three datasets with a total of 11617 micrograph stacks were individually collected in the same manner. These stacks were motion corrected with MotionCor2^37^ with a binning factor of 2, resulting in a pixel size of 1.07 Å, meanwhile dose weighting was performed. The defocus values were estimated using CTFFIND4^38^. For the ATP-bound ABCD1, 4302 micrograph stacks were individually collected with EPU software^39^ on a Titan Krios microscope at 300 kV equipped with a K3 Summit direct electron detector (Gatan) and a GIF Quantum energy filter (Gatan), at a nominal magnification of 29,000× with defocus values from −2.0 to −1.8 μm. For these stacks, motion correction and dose weighting were performed with patch motion correction with a Fourier cropping factor of 0.5, resulting in a pixel size of 1.07. Meanwhile, the defocus values were estimated using Patch CTF estimation^40^.

### Cryo-EM data processing

For the apo ABCD1 datasets, 2,194,963 and 1,899,981 particles were automatically picked from dataset 1 and dataset 2, using Relion 3.1^41,42^, respectively. After 2D classification, 1,174,746 particles from dataset 1 and 900,138 particles were selected and subjected to global search 3D classification. Then we merged particles from two datasets and after several rounds of 3D classification, 360,311 particles were selected for 3D auto-refinement. We adopted 3D auto-refinement of the particles with an adapted mask and yielded a reconstruction with an overall resolution of 3.53 Å, followed by CTF refinement and Bayesian Polishing. In the whole process, C1 symmetry was used and no other type of symmetries was applied.

For C22:0-CoA-bound ABCD1 datasets, 1,860,455 particles from Dataset 1, 2,583,043 particles from Dataset 2 and 2,500,479 particles from Dataset 3 were picked and subjected to 2D and initial global search 3D classification with a C1 symmetry, respectively. Then C2 symmetry was applied for further global 3D classification. 467,598 particles were selected from three datasets and imported into cryoSPARC3.2^40^ for ab-initio reconstruction and heterogeneous refinement. Ultimately, 336,741 particles were selected for further refinement and yielded reconstruction map at resolution of 3.59 Å with a C2 symmetry.

The procedures for ATP-bound ABCD1 were performed entirely on the cryo-SPARC3.2^40^. 1,878,754 particles were automatically picked from 4,302 micrographs and subjected to 2D classification, after which, 972,858 particles were selected and used to *ab-initial* reconstruction and heterogeneous refinement with a C1 symmetry. Then 550,864 particles were selected and applied to 4 rounds of ab-initial reconstruction and heterogeneous refinement with C2 symmetry being imposed to remove bad particles as much as possible. Ultimately, a reconstruction map at resolution of 2.79 with C2 symmetry were yielded by 528,274 Particles.

All resolutions of reconstruction map were estimated using the gold-standard Fourier shell correlation 0.143 criterion^43^.

### Model building and refinement

Since density for the segment residues from PMP-4 missed in all the three reconstruction maps, we numbered all the residues in these three structures corresponding to the sequence number of human ABCD1. The model of apo ABCD1 was manually rebuilt in COOT^44,45^ and refined using Real-space refinement in Phenix with secondary structure and geometry restraints^46^. The final model contains residues 68-337, 241-355, 371-436, and 436-725 in Chain A. For Chain B, residues including 68-337, 241-355, 371-435 and 463-720 were built according to the map. Two extra densities between TM5 and TM6 can be observed in each TMD, and we fit a 18:0 Lyso PE molecule in each density. For model of C22:0-CoA-bound ABCD1, residues 67-429 for TMDs, two C22:0-CoA and two CHS were built in the map manually. The density for the NBDs is too pool for us to build the structure accurately, so we fit the NBDs from ATP-bound ABCD1 and refine in Phenix. For the ATP-bound ABCD1, A homology outward-facing model of ABCD1 was generated by the SWISS-MODEL server, using the EM structure of human ABCD4 (PDB code 6jbj) as the reference. Residues 64-345, 383-435 for TMDs and 461-685 for NBDs were built in the map, and two prominent densities between two NBDs allowed us to fit two ATP/Mg^2+^.

The structural model was validated by Phenix and MolProbity^47,48^ and the model refinement and validation statistics were summarized in Extended Data Table 1. UCSF ChimeraX ^49^and PyMOL (https://pymol.org) were used for preparing the figures.

## Notes

### Competing Interest Statement

The authors have declared no competing interest.

