## Supplemental data for "Structural basis of substrate recognition and translocation by human ABCD1"

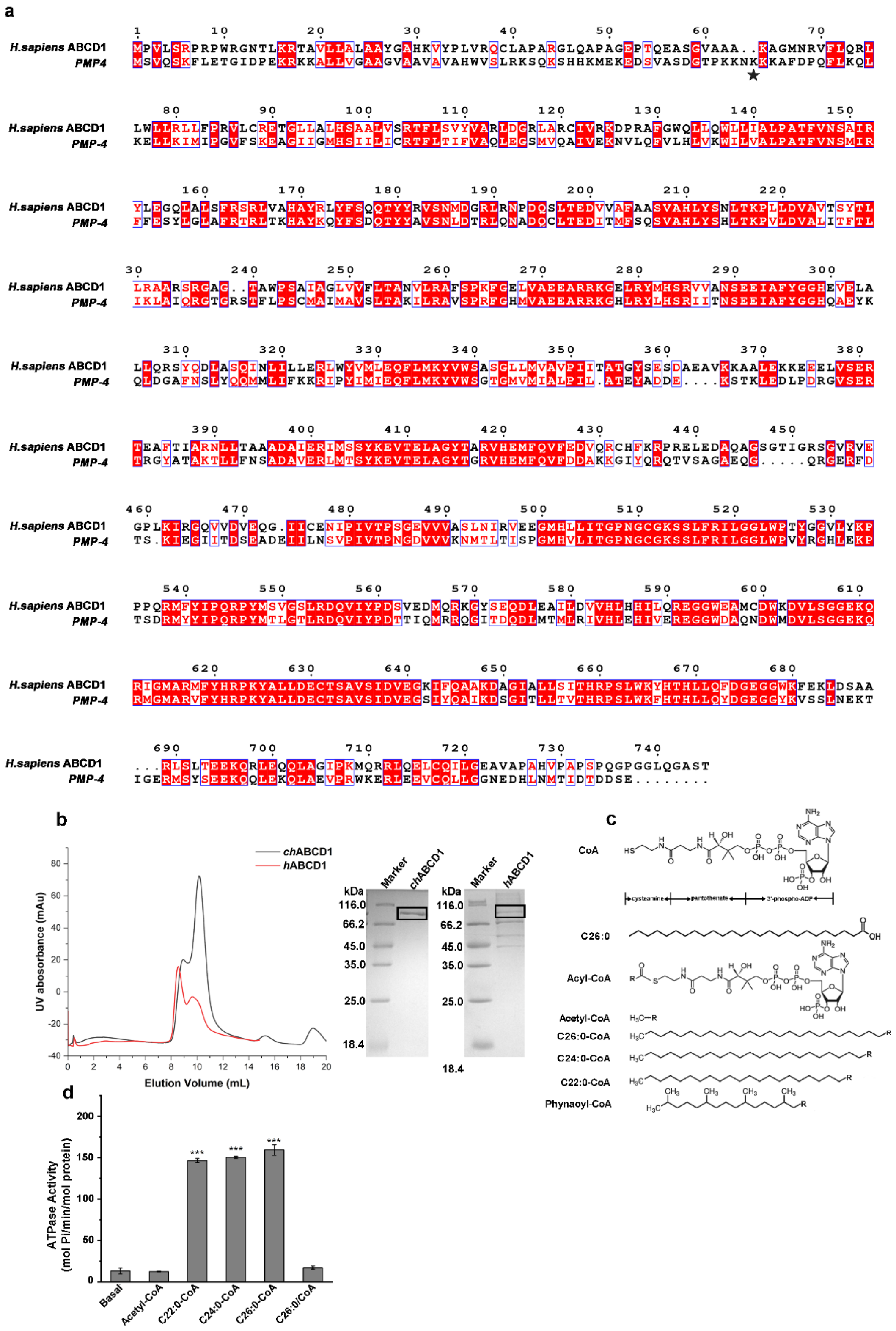


**Extended Data Fig. 1 | Purification and biochemical characterization of hABCD1 and chABCD1. a,** Sequence alignment of human ABCD1 and PMP-4. The substitution site for the chimeric ABCD1 construction is indicated by an asterisk. **b,** Size exclusion chromatography of hABCD1 and chABCD1. The peak fractions of 10 mL for hABCD1 and 10.2 mL for chABCD1 were pooled and concentrated for biochemical and structural studies. The purified protein was visualized by Coomassie-blue stained SDS-PAGE and bands for chABCD1 and hABCD1 were indicated by black rectangles. **c,** Structural formulas of potential substrates for ABCD1 used in the ATPase activity assays. **d,** Stimulated ATPase activity of human ABCD1 (hABCD1) over various CoA esters of very long chain fatty acid very long (VLCFA-CoAs) or acetyl-CoA. All data points analyzed above represent means of three independent measurements. Error bars indicate standard deviation. Unpaired two-sided t-test is used for the comparison of statistical significance. The P values of <0.05, 0.01, and 0.001 are indicated with *, ** and ***.

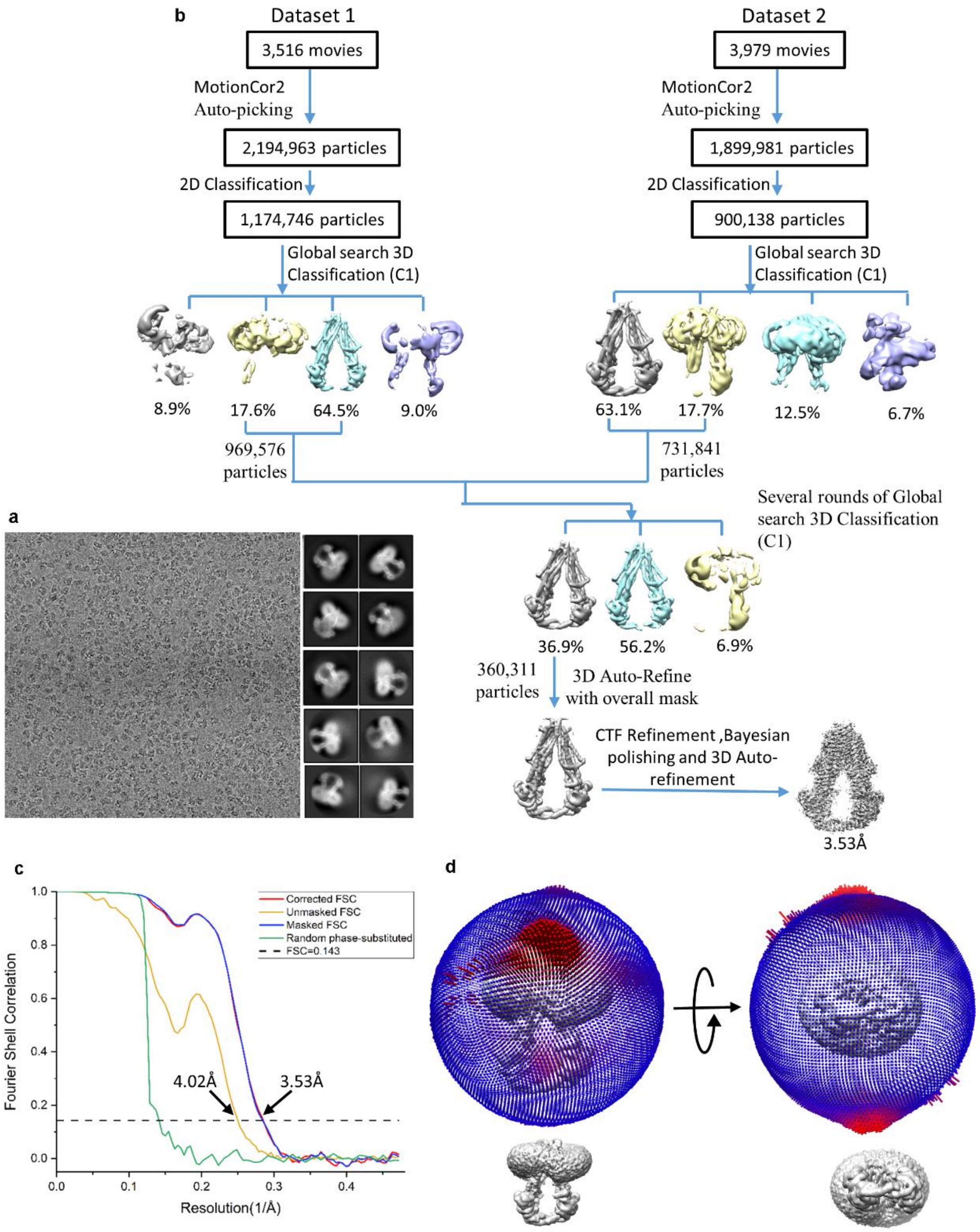
b
Dataset 1
3,516 movies
MotionCor2
Auto-picking
2,194,963 particles
2D Classification
1,174,746 particles
Global search 3D
Classification (C1)
8.9%
17.6%
64.5%
9.0%
969,576
particles
Dataset 2
3,979 movies
MotionCor2
Auto-picking
1,899,981 particles
2D Classification
900,138 particles
Global search 3D
Classification (C1)
63.1%
17.7%
12.5%
6.7%
731,841
particles
Several rounds of Global
search 3D Classification
(C1)
36.9%
56.2%
6.9%
360,311
particles
3D Auto-Refine
with overall mask
CTF Refinement ,Bayesian
polishing and 3D Auto-
refinement
3.53Å
a
c
Corrected FSC
Unmasked FSC
Masked FSC
Random phase-substituted
FSC=0.143
Fourier Shell Correlation
Resolution(1/Å)
4.02Å
3.53Å
1.0
0.8
0.6
0.4
0.2
0.0
0.0
0.1
0.2
0.3
0.4
d

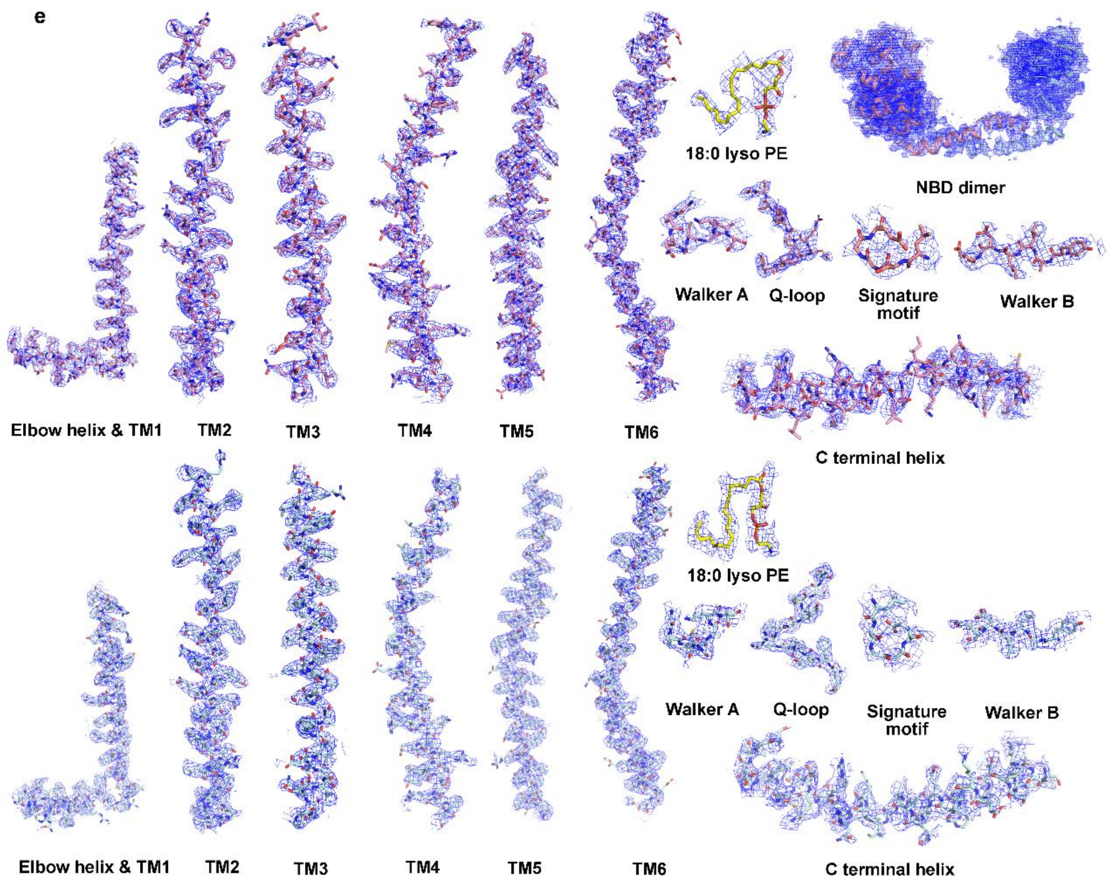


**Extended Data Fig. 2 | Cryo-EM analysis of apo-form ABCD1. a,** Representative cryo-EM micrographs and 2D averages. **b,** Flowchart for cryo-EM data processing. **c,** Gold-standard FSC curve for apo-form ABCD1 map generated using Relion 3.1. **d,** Euler angle distribution of the classified particles used for the final 3D refinement of the overall map. **E,** EM maps for representative segments of apo-form ABCD1. The structure was reconstructed with C1 symmetry, so all the represent segments in two monomers are presented. Contour levels are set at 5σ for TM1-6 and two 18:0 Lyso PE molecules, and at 4σ for the NBDs including Walker A, Q-loop, signature motif, and Walker B. The contour level for C-terminal coiled-coil helix is set at 3σ.

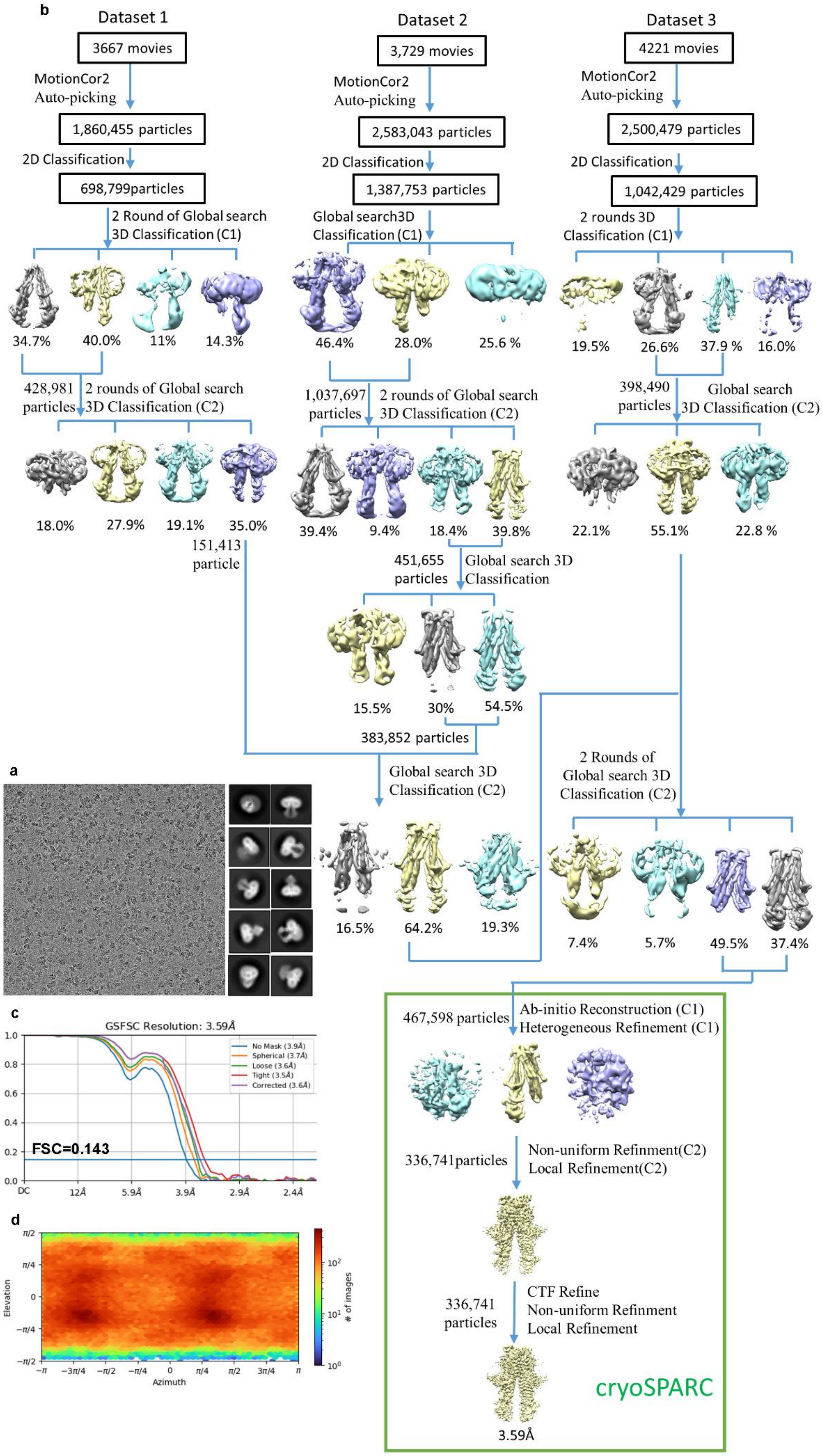
b
Dataset 1
3667 movies
MotionCor2
Auto-picking
1,860,455 particles
2D Classification
698,799particles
2 Round of Global search
3D Classification (C1)
34.7%
40.0%
11%
14.3%
428,981
particles
2 rounds of Global search
3D Classification (C2)
18.0%
27.9%
19.1%
35.0%
151,413
particle
Dataset 2
3,729 movies
MotionCor2
Auto-picking
2,583,043 particles
2D Classification
1,387,753 particles
Global search3D
Classification (C1)
46.4%
28.0%
25.6 %
1,037,697
particles
2 rounds of Global search
3D Classification (C2)
39.4%
9.4%
18.4%
39.8%
451,655
particles
Global search 3D
Classification
15.5%
30%
54.5%
383,852 particles
Global search 3D
Classification (C2)
16.5%
64.2%
19.3%
Dataset 3
4221 movies
MotionCor2
Auto-picking
2,500,479 particles
2D Classification
1,042,429 particles
2 rounds 3D
Classification (C1)
19.5%
26.6%
37.9 %
16.0%
398,490
particles
Global search
3D Classification (C2)
22.1%
55.1%
22.8 %
2 Rounds of
Global search 3D
Classification (C2)
7.4%
5.7%
49.5%
37.4%
467,598 particles
Ab-initio Reconstruction (C1)
Heterogeneous Refinement (C1)
336,741particles
Non-uniform Refinment(C2)
Local Refinement(C2)
336,741
particles
CTF Refine
Non-uniform Refinment
Local Refinement
cryoSPARC
3.59Å
a
c
GSFSC Resolution: 3.59Å
No Mask (3.9Å)
Spherical (3.7Å)
Loose (3.6Å)
Tight (3.5Å)
Corrected (3.6Å)
FSC=0.143
DC
12Å
5.9Å
3.9Å
2.9Å
2.4Å
d
Elevation
Azimuth
### of images

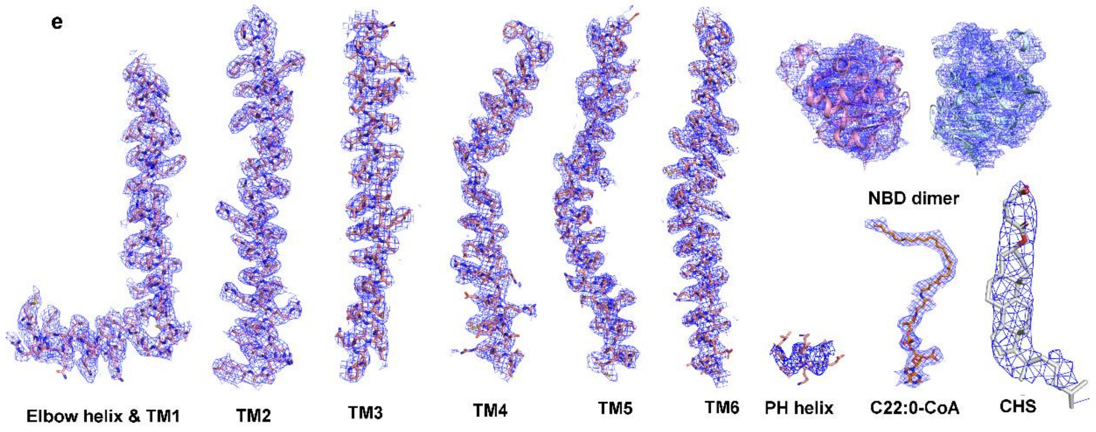


**Extended Data Fig. 3 | Cryo-EM analysis of C22:0-CoA-bound ABCD1. a,** Representative cryo-EM micrographs and 2D averages. **b,** Flowchart for cryo-EM data processing. **c**, Gold-standard FSC curve for C22:0-CoA-bound ABCD1 map generated using cryoSPARC 3.2. **d,** Euler angle distribution of the classified particles used for the final 3D refinement of the overall map**. e,** EM densities of representative segments of the structure of C22:0-CoA-bound ABCD1. The structure was reconstructed with C2 symmetry, and only one monomer is presented. Contour levels are set at 5σ for TM1-6, C22:0-CoA, and at 4σ for CHS. The contour level for the NBD is set at 3.5σ.

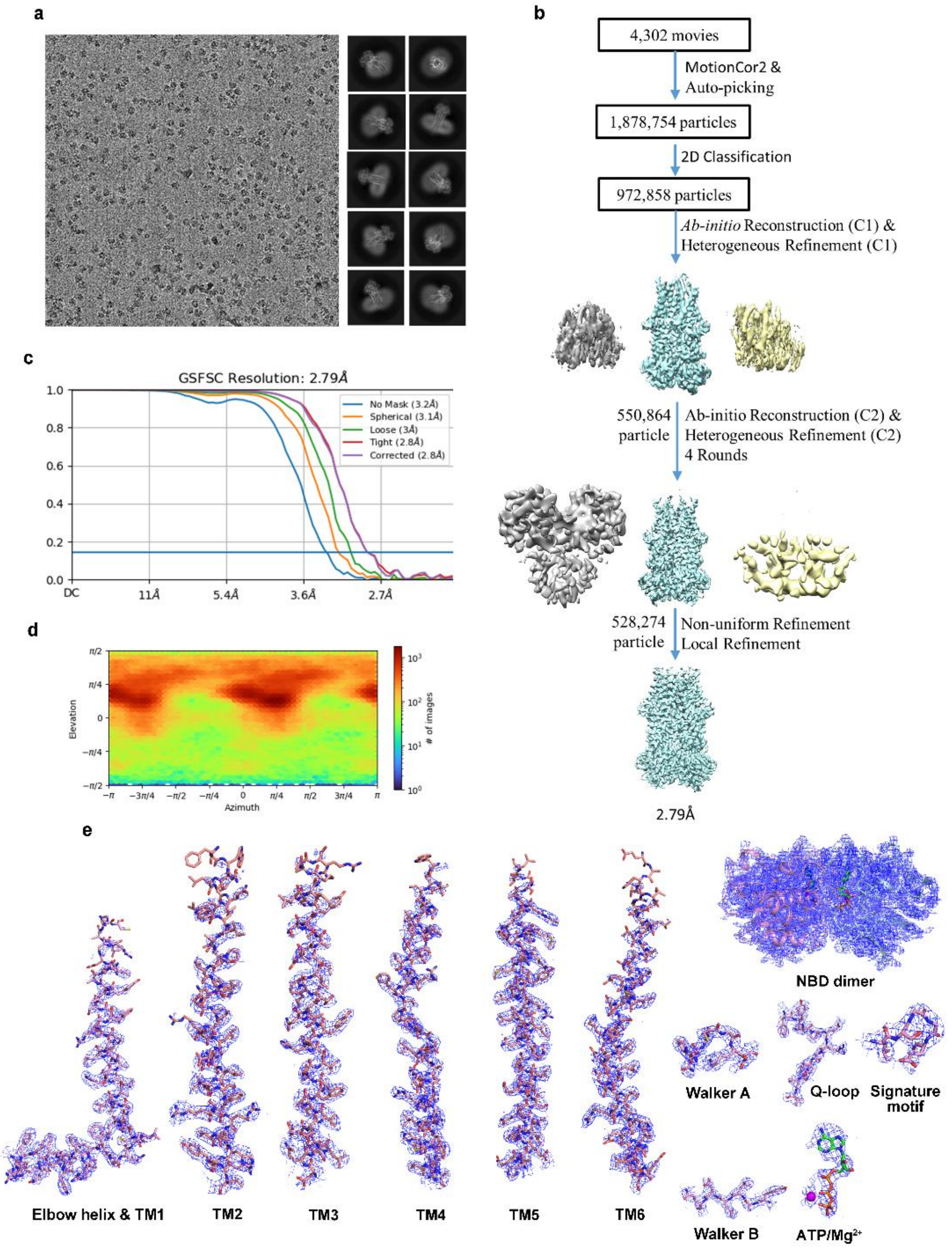


**Extended Data Fig. 4 | Cryo-EM analysis of ATP-bound ABCD1. a,** Representative cryo-EM micrographs and 2D averages. **b,** Flowchart for cryo-EM data processing. **c**, Gold-standard FSC curve for ATP-bound ABCD1 map generated using cryoSPARC 3.2. **d,** Euler angle distribution of the classified particles used for the final 3D refinement of the overall map**. e,** EM maps for representative segments of ATP-bound ABCD1. **e,** EM densities of representative segments of the structure of ATP-bound ABCD1. The structure was imposed with C2 symmetry, and segments only in one monomer are presented. Contour levels are set at 5σ for TM1-6, NBD and ATP/$Mg^{2+}$.

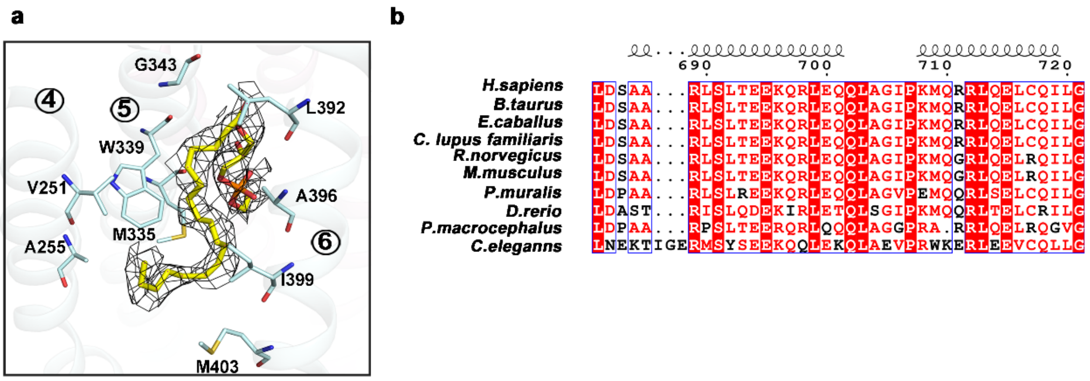


**Extended Data Fig. 5 | EM map for a lipid molecule and sequence alignment for the C-terminal helix. a,** a 18:0 Lyso PE molecule was fitted into the density (Contour level 5σ) between TM5 and TM6. **b,** Multiple-sequence alignment of the C-terminal helix among ABCD1 homologs.

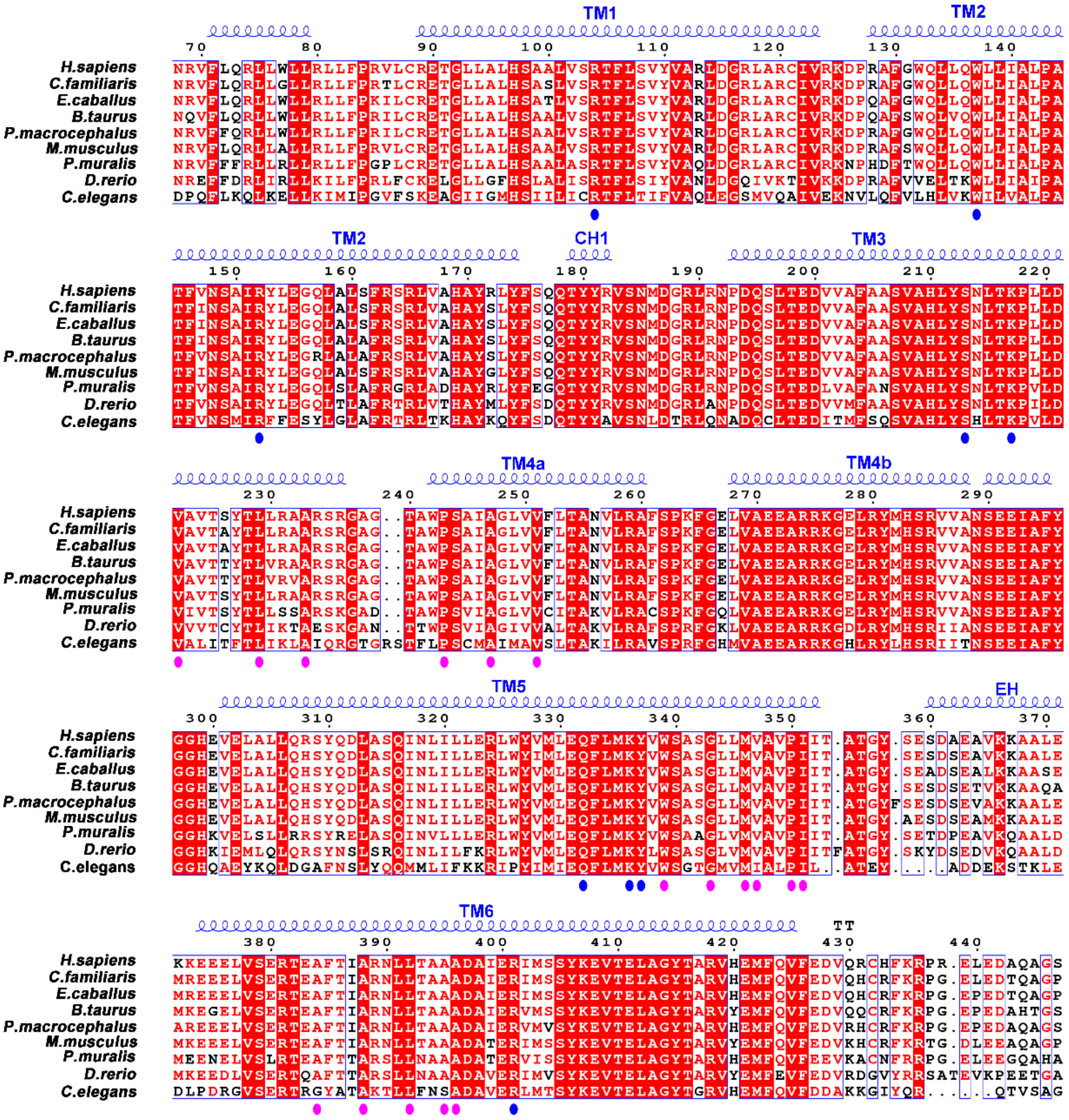

**Extended Data Fig. 6 | Sequence alignment of acyl-CoA binding sites among ABCD1 homologs.** Circles below the alignment indicate residues interacting with the CoA portion (blue) or the acyl chain (pink) portion of the substrate.

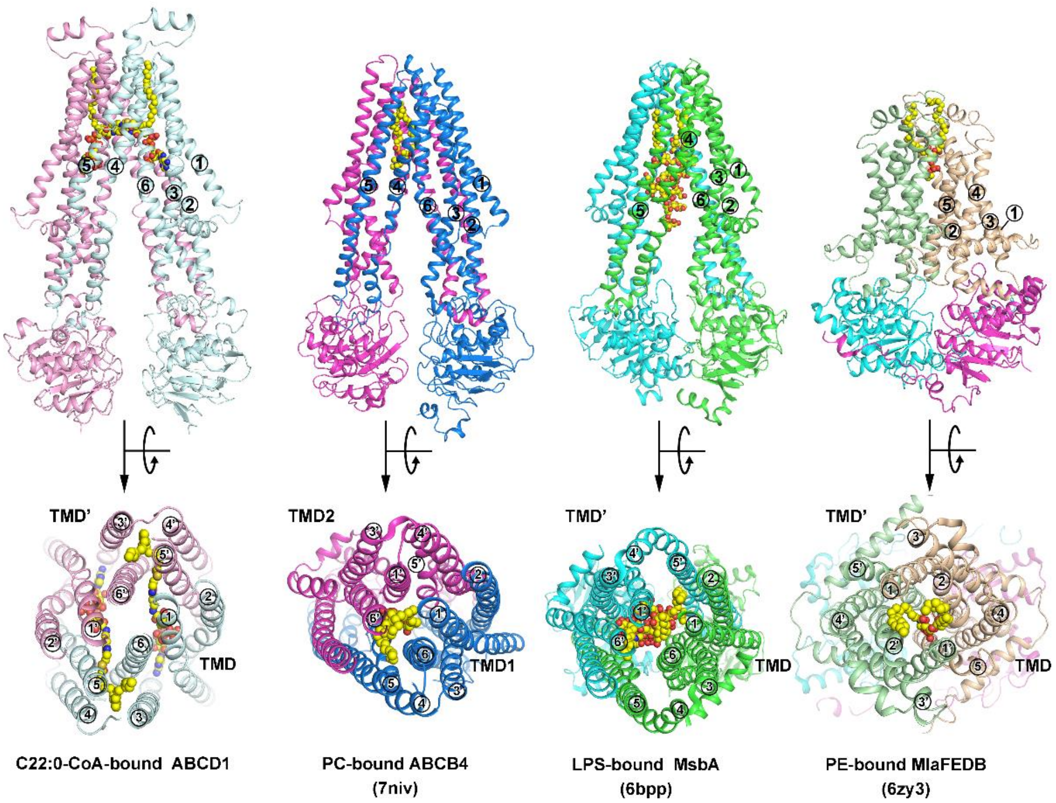


**Extended Data Fig. 7 | Substrate binding patterns adopted by known ABC transporters for amphipathic molecules with long acyl chains.** Substrates in these structures are displayed with spheres.

**Extended Data Table 1 | Cryo-EM data collection, refinement and validation statistics**

| | Apo ABCD1 (EMB-XXXX,PDB XXX) | C22:0-CoA-bound ABCD1 (EMB-XXXX,PDB XXX) | ATP-bound ABCD1 (EMB-XXXX,PDB XXX) |
|---|---|---|---|
| **Data collection and processing** | | | |
| Magnification | 22500 | 22500 | 81000 |
| Voltage (kV) | 300 | 300 | 300 |
| Camera | Gatan K3 summit | Gatan K3 summit | Gatan K3 summit |
| Electron exposure (e–/Å2) | 60 | 60 | 54 |
| Defocus range (μm) | -2.0 to -1.5 | -2.0 to -1.5 | -1.8 to 1.5 |
| Pixel size (Å) | 1.07 | 1.07 | 1.07 |
| Symmetry imposed | C1 | C2 | C2 |
| Initial particle images (no.) | 4,094,944 | 6,943,977 | 1,878,754 |
| Final particle images (no.) | 360,311 | 336,741 | 550,864 |
| Map resolution (Å) | 3.53 | 3.59 | 2.79 |
| FSC threshold | 0.143 | 0.143 | 0.143 |
| Map resolution range (Å) | 2.5-4.5 | 2.5-4.5 | 2.0-3.2 |
| **Refinement** | | | |
| Initial model used (PDB code) | Ab initio | Ab initial | 6jbj |
| Model resolution (Å) | 3.53 | 3.59 | 2.79 |
| FSC threshold | 0.143 | 0.143 | 0.143 |
| Map sharpening B factor ($Å^2$) | -117.5 | -185.8 | -116.8 |
| Model composition | | | |
| Nonhydrogen atoms | 9854 | 9463 | 9018 |
| Protein residues | 1223 | 1172 | 1120 |
| Ligands | 2 | 4 | 4 |
| Mean B factors (Å2) | | | |
| Protein | 61.77 | 89.73 | 39.77 |
| Ligand | 51.01 | 57.83 | 28.26 |
| R.m.s. deviations | | | |
| Bond lengths (Å) | 0.010 | 0.019 | 0.009 |
| Bond angles (°) | 0.975 | 1.850 | 0.992 |
| **Validation** | | | |
| MolProbity score | 2.68 | 2.75 | 1.97 |
| Clashscore | 15.68 | 22.01 | 7.69 |
| Poor rotamers (%) | 3.95 | 3.66 | 1.80 |
| Ramachandran plot | | | |
| Favored (%) | 90.64 | 92.10 | 94.95 |
| Allowed (%) | 8.95 | 7.73 | 5.05 |
| Disallowed (%) | 0.41 | 0.17 | 0.00 |
